# Identification and functional analysis of recent IS transposition events in rhizobia

**DOI:** 10.1101/2024.03.21.586147

**Authors:** Ezequiel G. Mogro, Walter O. Draghi, Antonio Lagares, Mauricio J. Lozano

## Abstract

Rhizobia are alpha- and betaproteobacteria that, through the establishment of symbiotic interactions with leguminous plants, are able to fix atmospheric nitrogen as ammonium. The successful establishment of a symbiotic interaction is highly dependent on the availability of nitrogen sources in the soil, and on the specific rhizobia strain. Insertion sequences (ISs) are simple transposable genetic elements that can move to different locations within the host genome and are known to play an important evolutionary role, contributing to genome plasticity by acting as recombination hot-spots, and disrupting coding and regulatory sequences. Disruption of coding sequences may have occurred either in a common ancestor of the species or more recently. By means of ISComapare, we identified Differentially Located ISs (DLIS) in nearly related rhizobial strains of the genera *Bradyrhizobium, Mesorhizobium, Rhizobium* and *Sinorhizobium*. Our results revealed that recent IS transposition events don’t seem to be playing a major role in adaptation. Nevertheless, DLIS could have a role enabling the activation and inactivation of certain genes that could dynamically affect the competition and survival of rhizobia in the rhizosphere.

## INTRODUCTION

Rhizobia are alpha- and betaproteobacteria that, through the establishment of symbiotic interactions with leguminous plants, are able to fix atmospheric nitrogen as ammonium. As a result of this interaction, rhizobia induce the development of a specialized root structure, the nodule, where they differentiate into bacteroids with the ability to fix N2, providing the plant with a source of nitrogen in exchange for photosynthetic carbon mostly in the form of dicarboxylic acids (Gourion et al., 2015; Jones et al., 2007; Oldroyd et al., 2011; Poole et al., 2018; Shumilina et al., 2023). These bacteria are thus of particular interest due to their potential to improve crop yields and reduce the need for synthetic nitrogen fertilizers (Herridge et al., 2008).

The successful establishment of a symbiotic interaction between rhizobia and legume plants is highly dependent on the availability of nitrogen sources in the soil, and on the specific rhizobia strain, being some strains highly competitive for the colonization of plant roots but presenting low N2-fixing efficiency (Checcucci et al., 2017). The colonization of the rhizosphere –the region of soil under the influence of plant roots– is the earliest step in the symbiotic process, and determines which rhizobia will end up in the nodules (Salas et al., 2017; Triplett and Sadowsky, 1992). The rhizosphere is a complex and dynamic environment where the interaction between plants, microorganisms, and soil particles takes place, thus playing a crucial role in plant growth and health (Walker et al., 2003). Nevertheless, the rhizosphere is not solely inhabited by rhizobia, but also by a diverse community of bacteria, fungi, and other microorganisms that compete for nutrients, but also for access to the host plant. The colonization of the rhizosphere is therefore a complex process that involves both plant-microbe interactions and competition with other rhizospheric organisms, and is influenced by various factors, including its growth rate, motility, and production of enzymes and other metabolites (Salas et al., 2017; Wheatley et al., 2020). Understanding the mechanisms and factors that influence rhizobia ability to colonize the rhizosphere can provide some insight into the dynamics of the rhizobia-legume symbiosis and potentially inform strategies for improving plant growth and productivity.

The genomes of rhizobia are very plastic, and are usually constituted by one chromosome, chromids –replicons of great size, difficult to distinguish from plasmids–, and several plasmids, some of them of great size –megaplasmids– and some smaller, generally considered as part of the accessory genome (Landeta et al., 2011; MacLean et al., 2007; Poole et al., 2018). Most of the genes required for the establishment of the symbiotic interaction with legume plants are located in such megaplasmids, or in certain cases, in chromosomal integrative and conjugative elements (ICEs) (Poole et al., 2018), and there is evidence suggesting that they are actively mobilizing within rhizobial populations (Wheatley et al., 2020). Furthermore, rhizobial genomes contain a great number of insertion sequences (IS) (Kaneko, 2002, 2000; Nelson et al., 2018, this work).

Insertion sequences (ISs) are simple transposable genetic elements that can move to different locations within the host genome (Siguier et al., 2014). They are known to play an important evolutionary role, contributing to genome plasticity by acting as recombination hot-spots, and disrupting coding and regulatory sequences (Consuegra et al., 2021; Iida et al., 2015; Siguier et al., 2014; Vandecraen et al., 2017). In rhizobia, genome architecture has been shown to be under constant modification, with replicons continuously being cointegrated and excised (Guo et al., 2003). Moreover, the role of tandemly repeated ISs as drivers of genomic recombination events was demonstrated in artificial evolution experiments (Arashida et al., 2021). ISs can also modify the expression of genes, being a known case in rhizobia the mutation of *Sinorhizobium meliloti* 1021 *expR* –a LuxR family transcriptional regulator that controls the expression of the symbiotically active exopolysaccharide (EPS) EPS II– by the insertion of a ISRm2011-1 insertion sequence (Pellock et al., 2002). However, the great amount of partial –most likely inactive– transposases suggest that some of these insertions could have taken place in a more distant common ancestor. The role of recent transposition events in rhizobia has not been thoroughly explored thus far.

In this study, we used ISCompare (Mogro et al., 2021) software to identify ISs that have changed their location in nearly related rhizobial strains, and analysed the genes disrupted by these differentially located ISs (DLISs). Through this study, we have gained a better understanding of the impact of recent IS transposition events on rhizobial lifestyle.

## MATERIALS AND METHODS

### Identification of differentially located ISs with ISCompare

ISCompare was used to compare the genomes of several rhizobial strains belonging to *Bradyrhizobium diazoefficiens, Bradyrhizobium japonicum, Mesorhizobium ciceri, Mesorhizobium loti, Rhizobium etli, Rhizobium leguminosarum*, and *Sinorhizobium meliloti*. Genome sequences were obtained from the National Center for Biotechnology Information (NCBI) assembly database (Kitts et al., 2016). The accession numbers for the sequences used in this study are compiled in Table S1.A.

The sequences were downloaded in FASTA, CDS FASTA, and GBFF format. The sequences used as IS database for ISCompare were downloaded from https://github.com/thanhleviet/Isfinder-sequences.

ISCompare (Mogro et al., 2021) is a tool used to find differential located ISs between two closely related strains. Here we made pairwise comparisons using a selected strain as reference and all the complete closed genomes available as of 5/Jan/2023. To account for the most confident results only the results classified as DLIS by ISCompare, and with a complete IS match, were analysed in this study. The software was run with the following parameters: E-value = 1e-10, minLength = 50, ISdiff = 50, scaffoldDiff = 20, minAlnLength = 50, surroundingLen = 500, surroundingLen2 = 500, shift = 0. The total number of DLIS located in accessory replicons –plasmids, megaplasmids– or in the main replicon –chromosome– was determined. DLIS counts were normalized by the species average size of the main and accessory replicons –the number of DLIS per 100,000 base pairs– to account for the difference in their sizes. The statistical analysis was done with python scripts using scipy (Virtanen et al., 2020). For the comparison of proportions Chi2 test was used summing the counts over all the strains within each species.

### Distribution of IS in rhizobia

A custom python script was used to count the genes and pseudogenes annotated as transposase or being part of an insertion sequence. In order to obtain this information, genbank files were processed with Biopython (Cock et al., 2009) and if the words ‘transposase’ or ‘insertion seq’ were found in a feature ‘product’ qualifier, a transposase was counted. In addition, the replicon information –transposon counts per replicon– was saved for further analysis.

### DLIS functional analysis

To determine whether an IS was located within an intergenic region that could correspond to a promoter, the following analysis was done. First, the two closest CDS to a DLIS insertion site were found using the “closest” command from bedtools software (Quinlan and Hall, 2010). When the insertion site was located within 150 nucleotides upstream of a CDS, the DLIS was considered to be interrupting a probable promoter region. To identify DLIS interrupting a probable operon, the two closest CDS to the insertion site were determined with bedtools “closest” command, and the orientation and a distance between these CDS was analysed. A possible operon was considered if the two CDS were in the same orientation, and separated by a maximum of 100 nucleotides.

In the case of DLIS inserted within coding sequences, a functional classification was made using COG (Cluster of orthologous groups database, Galperin et al., 2021). COGs were assigned to each gene using the eggNOG-mapper web server with the eggNOG 5 database (Huerta-Cepas et al., 2019).

Figures were plotted using python matplotlib (Hunter, 2007) and seaborn (Waskom, 2021). Whenever it was required, final figures were edited with Inkscape v 1.3 (The Inkscape team, https://inkscape.org/).

## RESULTS

### Analysis of IS and DLIS distribution in rhizobia

The genomes of all *B. diazoefficiens, B. japonicum, M. ciceri, M. loti, R. etli, R. leguminosarum*, and *S. meliloti* strains containing a complete assembly level were downloaded from NCBI Assembly database (Kitts et al., 2016, accessed 5/Jan/2023) and pairwise comparisons to identify DLIS were made with ISCompare (Table S1.B). For each species, a specific strain was used as reference for all pairwise comparisons. In order to estimate the proportion of active IS –DLIS with respect to the total number of IS– the total number of IS in a determined genome was estimated as the number of transposases plus the number of other genes annotated with functions related to insertion sequences (Figure 1, Table S1.C). Although the values are not exact –some IS may have been counted twice–, the estimated numbers of total ISs were in accordance with the previously reported (Kaneko, 2002, 2000; MacLean et al., 2007; Nelson et al., 2018). A recurrent observation is that chromosomal replicons tend to have lower number of total ISs whereas plasmids have a considerably higher amount (Figure 1.B, Table S1.E). This does not appear to be the case for DLIS (Figure 2.B., Table S2) which were detected with comparable frequency in plasmids (less but not significantly so) and chromosomal replicons. Of the rhizobia studied here, *Sinorhizobium* and *Bradyrhizobium* were the ones that showed the greatest number of DLIS (Figure 2.A) which agrees with the fact that they also present the highest number of ISs in their genomes (Figure 1.E-F, Table S1.C). Further, they also presented the highest proportion of IS that have recently changed their location in the genome, calculated as the number of DLIS / (total IS on ref strain + total IS on target strain) (Figure 3.A). Finally, a weak but significant correlation between the number of DLIS and the total number of IS was observed (Figure 3.B, R^2^ = 0.11; p-value = 4.55 e5).

**Figure 1.**
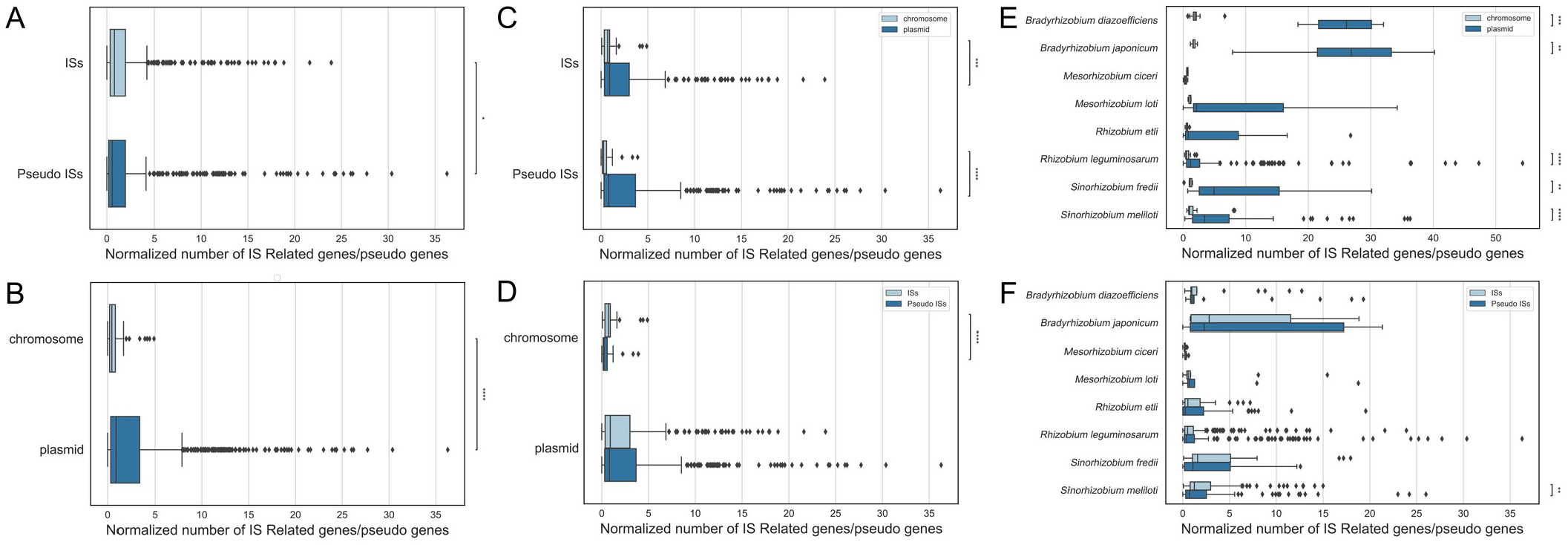
Distribution of transposase and IS related genes. A. Normalized IS and pseudo IS counts. B. Normalized number of IS elements by replicon. C-D. Nor malized numbers of ISs and pseudo ISs by replicon. E. Normalized number of IS elements by species and replicon. F. Normalized number of IS and pseudo ISs by species. The number of transposases and IS related genes was estimated from the genbank files annotation using custom python scripts that looked for the terms ‘transposase’ and ‘insertion seq’ in the product descriptions of coding sequences and pseudo genes. Normalized counts were calculated as ISs / 100,000 bp. Significance was determined using the Mann-Whitney test and the normalized counts (*: 1.00e-02 < p <= 5.00e-02; * *: 1.00e-03 < p <= 1.00e-02; * * *: 1.00e-04 < p <= 1.00e-03; ****: p <= 1.00e-04).

**Figure 2.**
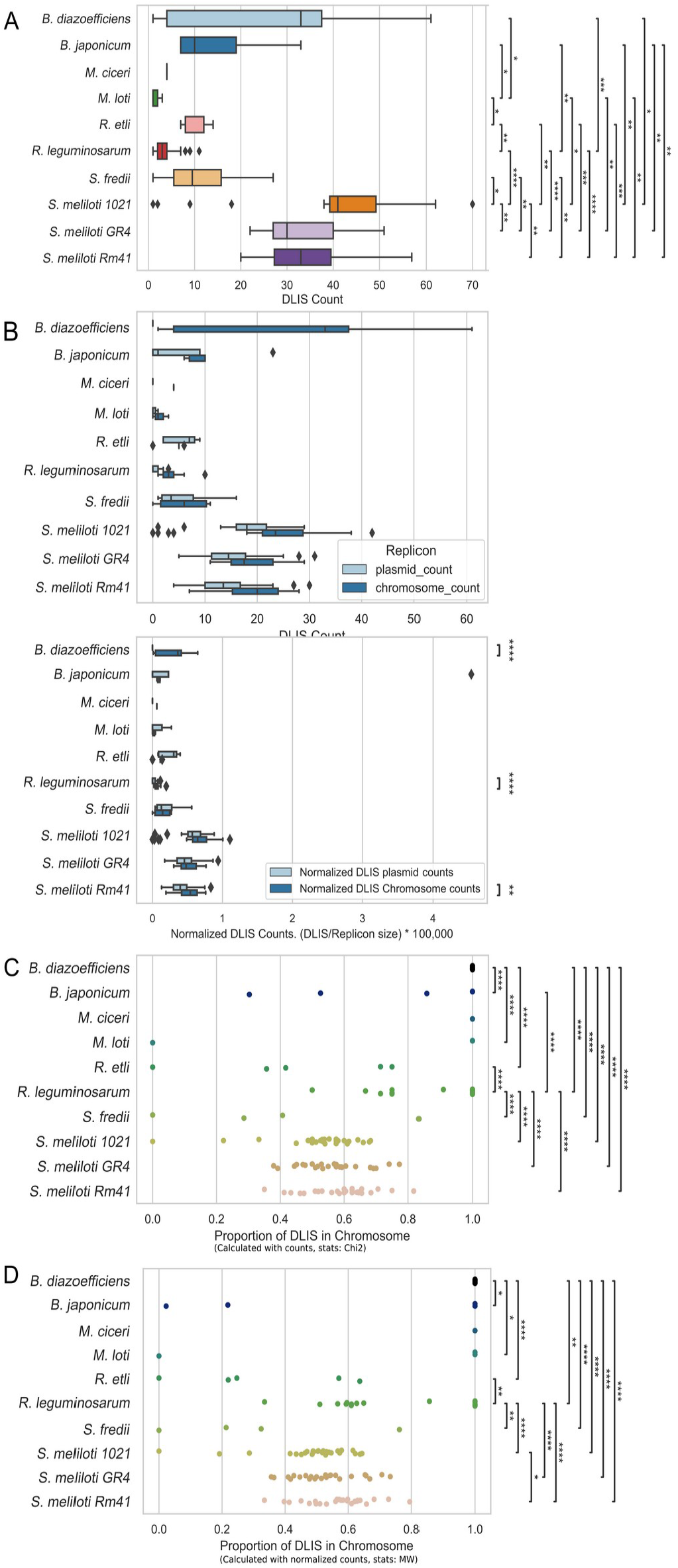
Distribution of DLIS. Chromosomes vs plasmids. A. Total DLIS counts. B. DLIS counts and normalized DLIS counts discriminated by rhizobia and replicon type. C. Proportions of DLIS in chromosomal replicons, calculated from DLIS counts. D. Proportions of DLIS in chromosomal replicons, calculated from DLIS normalized counts. DLIS were identified with ISCompare, and those with a full match for an insertion sequence and confidently identified as DLIS were analyzed. Significance was determined either using Chi2 test and the total DLIS counts in each species, or the Mann-Whitney test and the normalized DLIS counts (DLIS counts / 100,000 bp). *: 1.00e-02 < p <= 5.00e-02; ^**^: 1.00e-03 < p <= 1.00e-02; ^***^: 1.00e-04 < p <= 1.00e-03; ^****^: p <= 1.00e-04.

**Figure 3.**
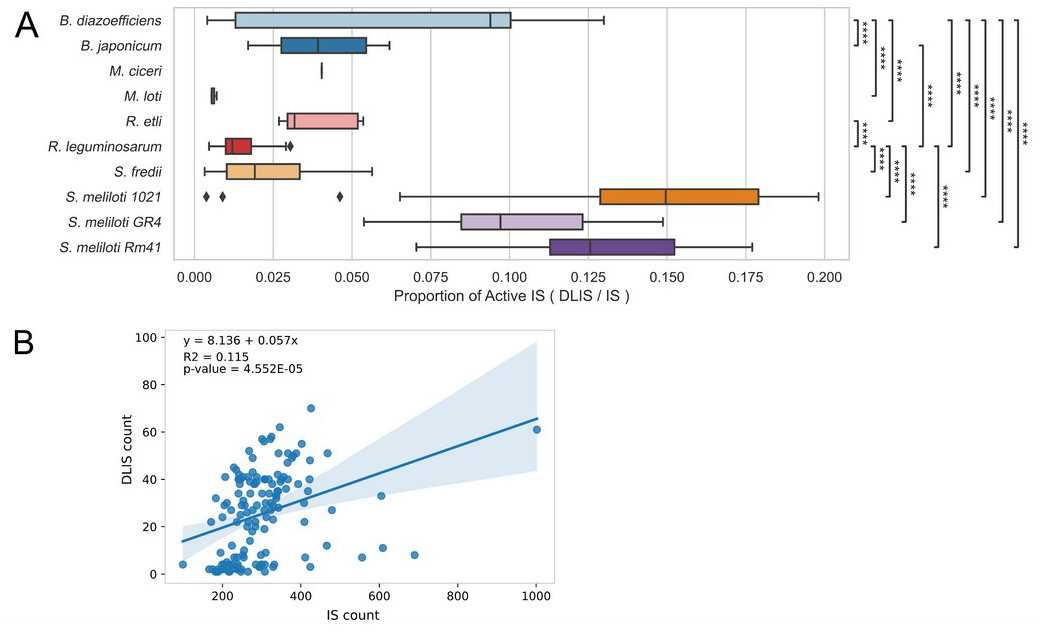
Proportion of recently active ISs. A. Proportion of active ISs. The proportion of recently active ISs was estimated as the relation of DLIS over the total IS count. Significance was determined using the Mann-Whitney test and the proportions of active ISs (DLIS / IS). *: 1.00e-02 < p <= 5.00e-02; ^**^: 1.00e-03 < p <= 1.00e-02; ^***^: 1.00e-04 < p <= 1.00e-03; ^****^: p <= 1.00e-04. B. Pearson correlation of IS counts and DLIS counts.

Next, the distribution of DLIS insertion sites within genes, intergenic regions and possible operons was determined. Most DLIS presented insertion sites in intergenic regions (ca. 60%, Figure 4.A, Table S3.A) and the lowest proportion was observed for DLIS inserted within putative operons. *Mesorhizobium* and *Rhizobium etli* were the only ones that presented a higher proportion of DLIS inserted within coding sequences (Figure 4.B-C), although they also presented the lowest numbers of DLIS. Remarkably, most of the intergenic DLIS appear not to interrupt promoter regions (Figure 4.C).

**Figure 4.**
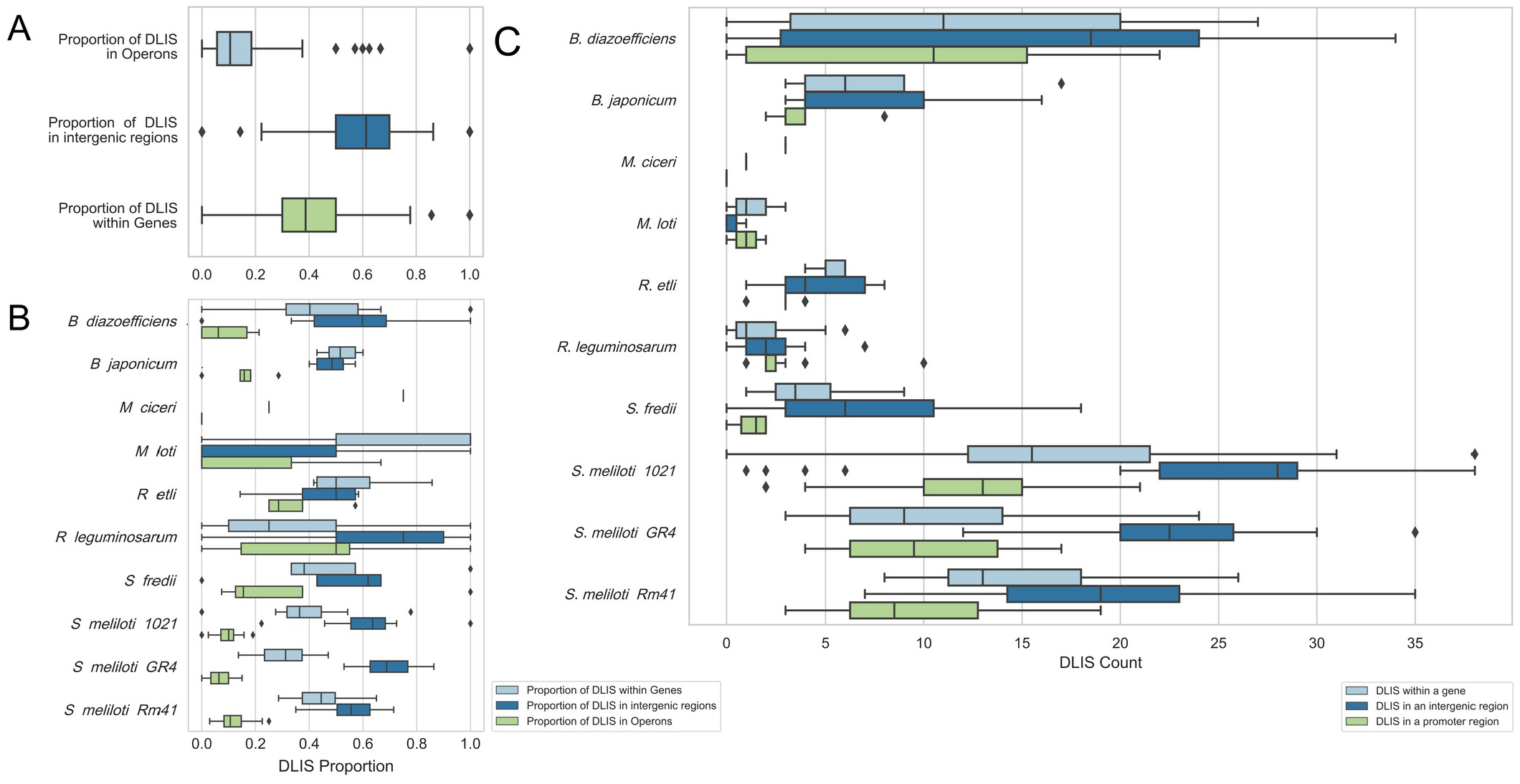
Distribution of DLIS in coding and intergenic regions. A. Proportion of DLIS inserted within genes, intergenic regions and putative operons. B. Proportion of DLIS inserted within genes, intergenic regions and putative operons discriminated by rhizobial species. C. Absolute count numbers of DLIS within genes, intergenic regions, and putative promoter regions. The count of the promoter regions is a subset of the counts for intergenic regions. When a DLIS was inserted within a short intergenic region between two divergent genes it was counted twice since it could be affecting the transcription of both genes. The location of the DLIS was determined using custom python scripts and the bedtools software as described in material and methods. Significance was determined using the Mann-Whitney test and the normalized counts.

### Functional distribution of genes interrupted by DLIS

To analyse the cellular functions affected by DLIS insertions, we recovered the amino acid sequences of the complete version of the proteins from the genomes that didn’t have the insertion sequence. All these genes were annotated with COGs using the eggNOG-mapper web server (Huerta-Cepas et al., 2019).

In the first analysis we used all the genes with an inserted DLIS (Figure 5). In such a case, each time a DLIS was inserted into a coding sequence of the genome used as reference, the disrupted gene could have been counted multiple times, as there could be many target genomes without the DLIS. In the second analysis, we only used the proteins with a DLIS insertion in the target genome, in that way, the complete sequence always corresponded to the reference genome which allowed us to avoid overcounting certain functions (Figure S1).

**Figure 5.**
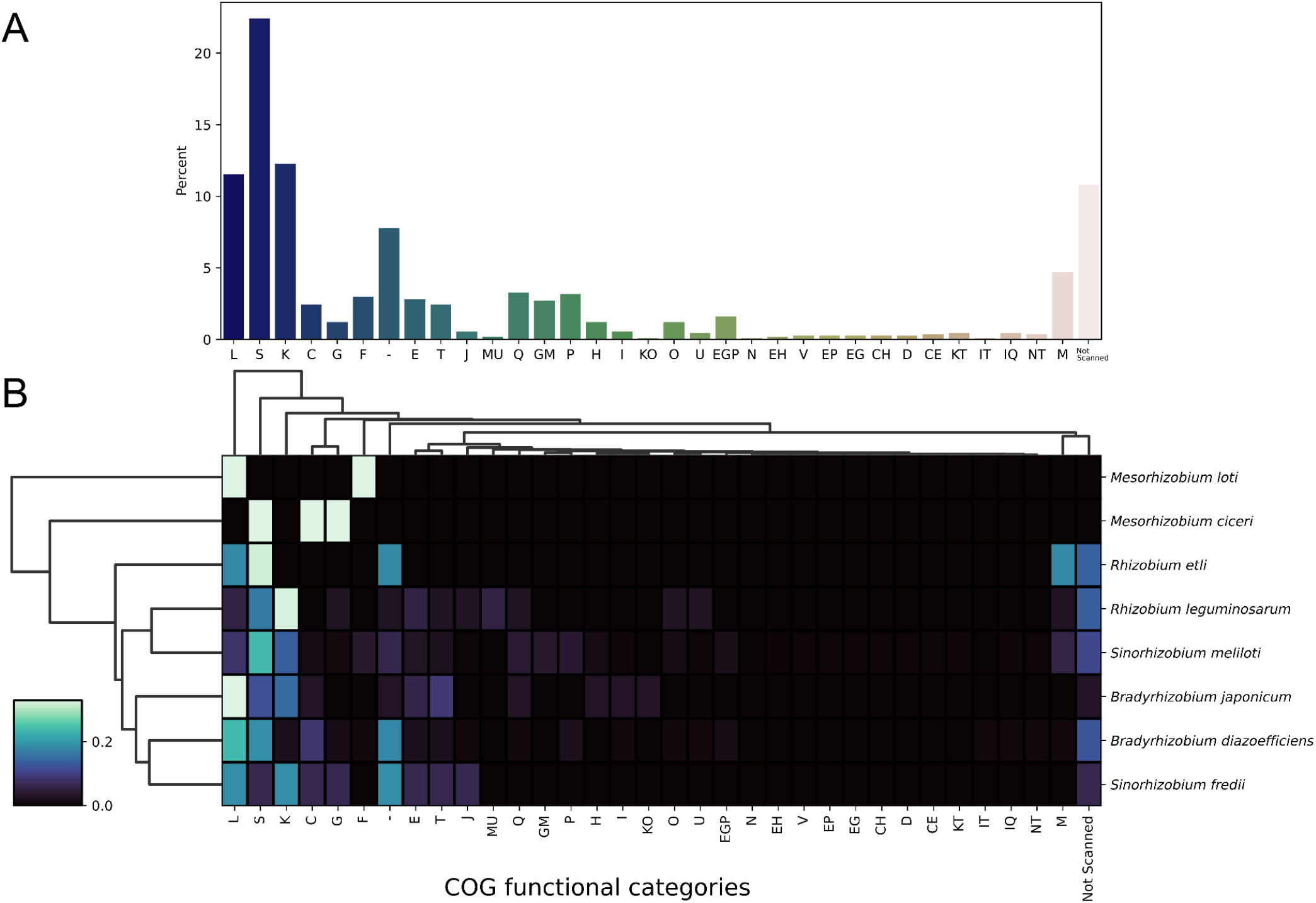
Distribution of COG functional categories. COGs functional categories for all the genes interrupted by a DLIS were assigned with eggNOG-mapper. A. Distribution of DLIS interrupted genes by COG functional category. B. The heatmap was generated with a custom python script using seaborn. COG categorías are as follows. -: Not assigned; A: RNA processing and modification; B: Chromatin Structure and dynamics; C: Energy production and conversion; D: Cell cycle control and mitosis; E: Amino Acid metabolism and transport, F: Nucleotide metabolism and transport, G: Carbohydrate metabolism and transport; H: Coenzyme metabolism; I: Lipid metabolism; J: Translation; K: Transcription; L: Replication and repair; M: Cell wall/membrane/envelope biogenesis; N: Cell motility; O: Post-translational modification, protein turnover, chaperone functions; P: Inorganic ion transport and metabolism; Q: Secondary metabolites biosynthesis, transport and catabolism; R: General Functional Prediction only; S: Function Unknown; T: Signal Transduction; U: Intracellular trafficking and secretion; V: Defense mechanisms: W: Extracellular structures: X: Mobilome: prophages, transposons; Y: Nuclear structure; Z: Cytoskeleton.

The overall results from the two analysis show that the top five COG categories –with the greatest number of DLIS– were S (Function unknown), K (Transcription), L (Replication, recombination and repair), M (Cell wall/membrane/envelope biogenesis) and Q (Secondary metabolites biosynthesis, transport and catabolism). L, S and K were also the principal categories in most rhizobia. Proteins with a classification in categories S and (-) corresponded majorly to hypothetical and conserved hypothetical proteins, or to proteins with unknown function. However, several proteins in the S category presented a more descriptive function, most remarkably some proteins with putative cellulose biosynthesis, chitinase, dihydroxyacetone kinase and nitrile hydratase activities. The L category contained DLIS that in most cases were inserted within genes encoding for transposases, integrases and reverse transcriptases, with only a few DLIS inserted in genes annotated as involved in DNA repair, nucleases, DNA polymerases, recombinases and methylases. For the K category, most of the proteins interrupted by DLIS corresponded to different families of transcriptional regulators, acetyltransferase domain containing proteins, and proteins involved in plasmid partition. Particularly, NodD1 and NifA presented DLIS insertions in *B. japonicum* USDA 6 and *S. meliloti* AK76 respectively. Of interest to us was that several proteins in the M (Cell wall/ membrane/envelope biogenesis) COG category were interrupted by DLIS.

In particular, a capsular polysaccharide biosynthesis export transmembrane protein, and a choline-glycine betaine transporter were mutated in *S. meliloti* 1021 and *S. meliloti* GR4 respectively. Also *lpxK*, involved in the modification (phosphorylation) of lipid IVa, was interrupted in *S. meliloti* GR4 and 1021 genomes. In the E category, most of the insertions were within different enzymes annotated as arginine/lysine/ornithine decarboxylase, aspartate/tyrosine/aromatic aminotransferase, choline dehydrogenase, ethanolamine ammonia lyase, and phosphoribosyl anthranilate isomerase (Usg). Other insertions that could be relevant were observed in proteins belonging to the categories T (Signal Transduction) and P (Inorganic ion transport and metabolism). In particular DLIS insertions were observed in adenylate/guanylate cyclases, in two component systems (both in Histidine kinases and response regulators), and in several Na^+^/H^+^, Mg^2+^/ Co^2+^, Fe^3+^ and K^+^ transport proteins. In the G category (Carbohydrate metabolism and transport) DLIS were found mainly in ABC transporters including a ribose/xylose/arabinose/galactoside and Branched-chain amino acid transport systems. Finally in the C category (Energy production and conversion), proteins annotated as aldehyde dehydrogenase, nitrite reductase, cytochrome bd terminal oxidase, and pyruvate ferredoxin/flavodoxin oxidoreductase activities presented DLIS insertions.

Certain genes presented DLIS insertions that were overcounted because they were mutated in the reference genome. These included genes annotated as major facilitator superfamily (MFS) transporter, hypothetical protein with adenylate kinase activity, polysaccharide biosynthesis protein (*lpsB2*), GNAT family N-acetyltransferase, LuxR family transcriptional regulator, Lrp/AsnC ligand binding domain-containing protein, LysR family transcriptional regulator, magnesium and cobalt transport protein CorA, methyltransferase domain-containing protein (annotated as NodS by eggnog mapper), dihydroxyacetone kinase (Dak2), DedA family protein, Nitrile hydratase, RES domaincontaining protein, tripartite tricarboxylate transporter, VIT family protein and the previously mentioned *lpxK*.

## DISCUSSION

Rhizobial plasmids are known to have a greater number of ISs than chromosomes and it has been shown that they can act as mediators of homologous recombination leading to genomic rearrangements (Guo et al., 2003). Surprisingly, in the case of DLIS, a greater number was observed in chromosomal replicons (Figure 2.C-D). DLIS are ISs inserted into highly conserved regions in a genome of a pair of closely related bacterial genomes and could therefore be broadly considered as active ISs. Our results appear to indicate that ISs tend to be more active in chromosomal replicons than in plasmids. This result was not expected, and could be a consequence of the high proportion of pseudo IS elements in plasmids (Figure 1.D), which, although perfectly capable of mediating homologous recombination, are inactive. To confidently identify recent transposition events, in this work we only considered DLIS with complete blast hits to ISs in the library. However, ISCompare also identifies DLISs involving partial IS blast hits which could be related to more ancestral insertions and were not considered.

Of the analysed rhizobia genera, *Sinorhizobium* and *Bradyrhizobium* presented the higher proportion of ISs, DLISs and active ISs. Furthermore, we found a weak but significant correlation (r=0.115, pvalue=4.552x10^-5^, Figure 3.B) between the number of ISs and the number of DLISs. Thus, a more important contribution of ISs to environmental adaptation could be expected for genomes with greater numbers of ISs. However, most rhizobial DLISs were found to be inserted within intergenic regions, and far from the 5’-ends of coding sequences (Figure 4), suggesting that ISs might play a more important role in genome plasticity and adaptation by acting as recombination hot-spots than through transposition per se. This interpretation has to be taken with care, since intergenic regions could encode for small ORFs and RNAs not yet annotated. Furthermore, only a low proportion of the identified DLIS are expected to generate polar mutations by interrupting putative operons (Figure 4.A-B).

Regarding the interrupted genes, the majority corresponded to COG categories L (mostly transposases), S (unknown function) and (not assigned), which reinforces that, in rhizobia, the adaptive mutations produced by IS transposition may be of less importance than those produced by IS-mediated homologous recombination. Nevertheless, we were able to identify some DLIS inserted within genes with a more precise annotation (Table 1). Some of those genes were reported to have a role in the symbiotic interaction with the plant host. Three of them corresponded to enzymes involved in the biosynthesis of the lipopolysaccharide (LPS), *lpxK, lpsB2 and kdsD*. LpxK catalyses the sixth step in the lipid A synthesis (Emptage et al., 2013); while LpsB2 is required for O-antigen biosynthesis and its mutants presented reduced motility, grew faster than the parental strain, and were more sensitive to maize benzoxazinones and polymyxin B (García-de los Santos and Brom, 1997; Ormeno-Orrillo et al., 2008). KdsD is required for the biosynthesis of 3-deoxy-d-manno-octulosonic acid (Kdo), a key sugar in the core region of LPS (Jenkins et al., 2023).

**Table 1.**
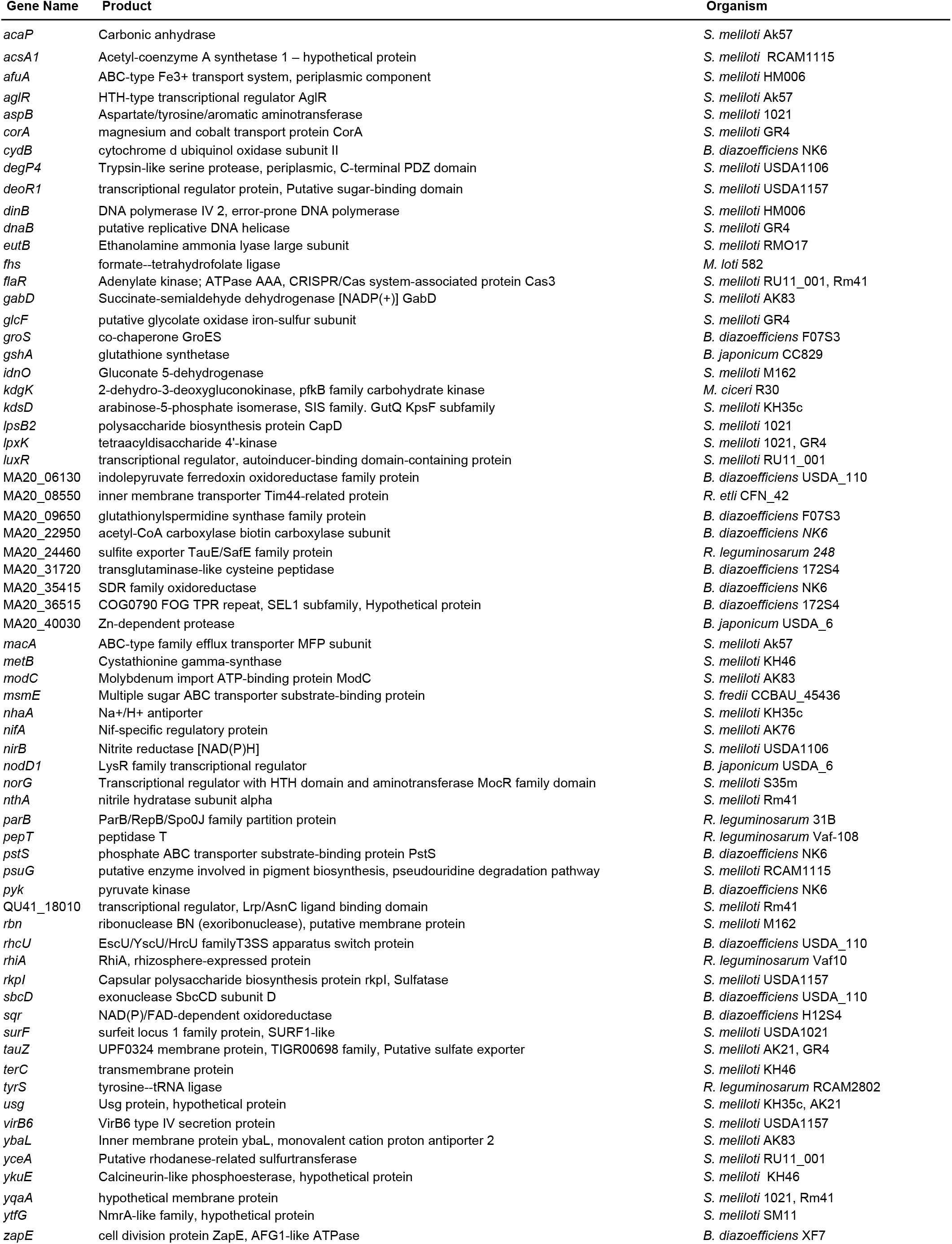
List of genes interrupted by a DLIS and annotated with a gene name in the reference strain.

Furthermore, a DLIS inserted within *rpkI*, a gene encoding for a protein involved in capsular polysaccharide biosynthesis, was found in *S. meliloti* USDA1157. *rhiA*, the first gene of the rhizosphere-expressed genes operon (*rhiABC*), that was reported to influence nodulation (Cubo et al., 1992), was found to be interrupted by a DLIS in *R. leguminosarum* Vaf10 strain.

The *nifA* gene of *S. meliloti* –a regulatory nitrogen fixation gene required for the induction of several key *nif* and *fix* genes (Agron et al., 1993)– presented an inserted DLIS on strain AK76. NifA null mutants induce white nodules in the roots of the host plant, have reduced swarming ability, and present lower levels acyl-homoserine lactones and of extracellular proteins (Gong et al., 2007). In *B. japonicum* USDA 6, NodD1, the positive regulatory protein of the *nodYABC* operon (Stacey, 1995), was found to be interrupted by a DLIS. In that case, a shorter ORF is annotated in the genome, and it could be possible that its product is still functional.

Another protein that was found to be interrupted by a DLIS is NirB. NirB, was shown to be involved in the nitrate assimilatory pathway, and to participate indirectly in NO synthesis, possibly contributing with the denitrification pathway (Ruiz et al., 2019), and thus, to the generation of greenhouse gases. GshA (glutathione synthetase) involved in glutathione biosynthesis, was interrupted in *B. japonicum* CC829. Glutathione has been shown to be important for growth under defined conditions, and to play an important role in symbiosis (Harrison et al., 2005; Sobrevals et al., 2006; Taté et al., 2012). Finally, Carbonic anhydrase (CA) –an enzyme involved in the interconversion of carbon dioxide and bicarbonate– was found to be interrupted in *S. meliloti* AK57. Although CA was shown not to be essential for nodule development and nitrogen fixation, it was hypothesized that it could help in the protonation of extracellular NH3 facilitating its diffusion and transport to the plant tissues (Kalloniati et al., 2009) .

Other genes encoding proteins not directly implicated in the interaction of rhizobia with their plant host, but that could have a role in rhizosphere colonization, were also found to be interrupted by DLISs, including *aspB* (aromatic aminotransferase), *acsA1* (Acyl-coenzyme A synthetases), *nthA* (nitrile hydratase subunit alpha), *cydB* (cytochrome d ubiquinol oxidase subunit II), *surF* (SURF1-like protein), *fhs* (formate-tetrahydrofolate ligase), *gabD* (Succinate-semialdehyde dehydrogenase), GlcF (glycolate oxidase complex, iron-sulphur subunit), *idnO* (Gluconate 5-dehydrogenase, GA5DH), metB (Cystathionine gamma-synthase), pyk (pyruvate kinase), and *eutB* (ethanolamine ammonialyase, large subunit). AspB, catalyses the transfer of an alpha amino group from aromatic amino acids, but also from aspartate, to different substrates. It was reported that it contributed, under high levels of exogenous tryptophan, to the biosynthesis of indole-acetic acid (IAA), and to nitrogen scavenging under nitrogen deprivation (Kittell et al., 1989). Also related to the IAA metabolism, *nthA* (coding for nitrile hydratase subunit alpha) was found interrupted by a DLIS in *S. meliloti* RM41 (Liu et al., 2017). An insertional mutation in *acsA1* was found in *S. meliloti* strain RCAM1115 which might impede its growth using acetate as carbon source (Aneja et al., 2002). GabD *–*Succinate-semialdehyde dehydrogenase– homologs are required for growth on γ-aminobutyrate (GABA) as the sole nitrogen source (Prell et al., 2009). GA5DH catalyses the transformation of 5-keto-d-gluconate to d-gluconate – that can enter the Entner-Doudoroff pathway to provide a source of carbon and NADP^+^– when energy is exhausted. However, under conditions of energy abundance, this enzyme catalyses the oxidation of dgluconate to 5-keto-d-gluconate which can be used as a transient means of energy storage (Zhang et al., 2009).

In the transport systems category several genes presented DLIS insertions in specific strains which could impair the absorption of nutrients, and thus their ability to compete for rhizosphere colonization. Among them, *afuA* (ABC-type Fe3+ transport system, periplasmic component), *corA* (magnesium and cobalt transport protein), *MA20_24460* (sulfite exporter TauE/SafE family protein), *tauZ* (putative sulfate exporter) *macA* (RND family efflux transporter MFP subunit), *modC* (Molybdenum import ATP-binding protein ModC), *rhcU* (type III secretion system export apparatus switch protein), *msmE* (Multiple sugar ABC transporter substrate-binding protein), *nhaA* (Na+/H+ antiporter), ybaL (monovalent cation proton antiporter 2) and *pstS* (phosphate ABC transporter substrate-binding protein PstS). Of these, CorA was shown to be important for growth on glucose at 21% O2 (Wheatley et al., 2017). MacA is the membrane fusion protein of a ABC-Type efflux transporter, a type 1 secretion system that transports diverse molecules (antibiotics and peptides) across the inner and outer membranes (Alav et al., 2021; Eda et al., 2011). ModC is required for nitrogen fixation on limiting levels of molybdate and was reported to be the high-affinity molybdate transporter in *B. diazoefficiens (Cheng et al*., *2016; Tresierra-Ayala et al*., *2011)*.

In the case of NhaA, it was shown to be induced under acid conditions, and it is likely that it functions by expelling protons toward the periplasm (Guerrero-Castro et al., 2018). Last, *pstS* is a part of the *pstSCAB* phosphate-specific transport operon that functions as highaffinity phosphate transporter (Guerrero-Castro et al., 2018). Moreover, an homologous of *Smc03123* (putative transcriptional regulator), *a* gene from *S. meliloti* 2011 which mutant was shown to have an altered fitness in the competence for rhizosphere colonization at 3 days post-inoculation (Salas et al., 2017), exhibited a DLIS in *S. meliloti* USDA1157.

Competition in the rhizosphere requires rhizobia to be able to use diverse compounds as carbon and nitrogen sources. Possessing a wide repertoire of metabolic pathways and transporters, enable rhizobia to thrive and compete for the nutrients exuded to the rhizosphere by the plant host. Dynamic inactivation of genes by DLIS could play an important role, granting a faster growth capability. However, it could also be detrimental, affecting posterior steps that occur during the interaction with the plant host, or under changing environmental conditions.

## CONCLUSION

In this work we analysed recent transposition events in rhizobia by identifying the DLISs by means of ISCompare. Our results revealed that recent IS transposition events don’t seem to be playing a major role in adaptation. This hypothesis is supported by the vast majority of DLIS being inserted within intergenic regions, and by the fact that most DLIS inserted within genes correspond to genes coding for hypothetical proteins. A more important role of ISs could be to mediate homologous recombination, allowing the dynamic interchange of information between chromosomal and plasmid replicons, and between different rhizobial species. Nevertheless, DLIS could have a role enabling the activation and inactivation of certain genes that could dynamically affect the competition and survival in the rhizosphere. To further characterize the role of DLIS in rhizobia, experimental evolution of single rhizobial isolates should be carried on. Through this type of experiment a clearer picture of the dynamics of IS transposition and recombination would be achieved.

## REFERENCES

Agron, P.G., Ditta, G.S., Helinski, D.R., 1993. Oxygen regulation of nifA transcription in vitro. Proc. Natl. Acad. Sci. 90, 3506–3510.10.1073/pnas.90.8.3506

Alav, I., Kobylka, J., Kuth, M.S., Pos, K.M., Picard, M., Blair, J.M.A., Bavro, V.N., 2021. Structure, Assembly, and Function of Tripartite Efflux and Type 1 Secretion Systems in Gram-Negative Bacteria. Chem. Rev. 121, 5479–5596.10.1021/acs.chemrev.1c00055

Aneja, P., Dziak, R., Cai, G.-Q., Charles, T.C., 2002. Identification of an Acetoacetyl Coenzyme A Synthetase-Dependent Pathway for Utilization of <scp>l</scp>-(+)-3-Hydroxybutyrate in Sinorhizobium meliloti. J. Bacteriol. 184, 1571–1577.10.1128/JB.184.6.1571-1577.2002

Arashida, H., Odake, H., Sugawara, M., Noda, R., Kakizaki, K., Ohkubo, S., Mitsui, H., Sato, S., Minamisawa, K., 2021. Evolution of rhizobial symbiosis islands through insertion sequence-mediated deletion and duplication 16.10.1038/s41396-021-01035-4

Checcucci, A., DiCenzo, G.C., Bazzicalupo, M., Mengoni, A., 2017. Trade, Diplomacy, and Warfare: The Quest for Elite Rhizobia Inoculant Strains. Front. Microbiol. 8, 2207.10.3389/FMICB.2017.02207

Cheng, G., Karunakaran, R., East, A.K., Poole, P.S., 2016. Multiplicity of Sulfate and Molybdate Transporters and Their Role in Nitrogen Fixation in Rhizobium leguminosarum bv. viciae Rlv3841. Mol. Plant-Microbe Interact. 29, 143–152.10.1094/MPMI-09-15-0215-R

Cock, P.J.A., Antao, T., Chang, J.T., Chapman, B.A., Cox, C.J., Dalke, A., Friedberg, I., Hamelryck, T., Kauff, F., Wilczynski, B., De Hoon, M.J.L., 2009. Biopython: Freely available Python tools for computational molecular biology and bioinformatics. Bioinformatics Bioinformatics, 1422–1423.10.1093/bioinformatics/btp163

Consuegra, J., Gaffé, J., Lenski, R.E., Hindré, T., Barrick, J.E., Tenaillon, O., Schneider, D., 2021. Insertion-sequence-mediated mutations both promote and constrain evolvability during a long-term experiment with bacteria. Nat. Commun. 12, 1–12.10.1038/s41467-021-21210-7

Cubo, M.T., Economou, A., Murphy, G., Johnston, A.W., Downie, J.A., 1992. Molecular characterization and regulation of the rhizosphere-expressed genes rhiABCR that can influence nodulation by Rhizobium leguminosarum biovar viciae. J. Bacteriol. 174, 4026– 4035.10.1128/jb.174.12.4026-4035.1992

Eda, S., Mitsui, H., Minamisawa, K., 2011. Involvement of the smeAB multidrug efflux pump in resistance to plant antimicrobials and contribution to nodulation competitiveness in Sinorhizobium meliloti. Appl. Environ. Microbiol. 77, 2855–2862.10.1128/AEM.02858-10

Emptage, R.P., Pemble, C.W., York, J.D., Raetz, C.R.H., Zhou, P., 2013. Mechanistic characterization of the tetraacyldisaccharide-1-phosphate 4′-kinase LpxK involved in lipid a biosynthesis. Biochemistry Biochemistry, 2280–2290.10.1021/BI400097Z/SUPPL_FILE/BI400097Z_SI_001.PDF

Galperin, M.Y., Wolf, Y.I., Makarova, K.S., Alvarez, R.V., Landsman, D., Koonin, E. V., 2021. COG database update: focus on microbial diversity, model organisms, and widespread pathogens. Nucleic Acids Res. 49, D274–D281.10.1093/NAR/GKAA1018

García-de los Santos, A., Brom, S., 1997. Characterization of Two Plasmid-borne lps β Loci of Rhizobium etli Required for Lipopolysaccharide Synthesis and for Optimal Interaction with Plants. Mol. Plant-Microbe Interact. 10, 891–902.10.1094/MPMI.1997.10.7.891

Gong, Z., Zhu, J., Yu, G., Zou, H., 2007. Disruption of nifA Gene Influences Multiple Cellular Processes in Sinorhizobium meliloti. J. Genet. Genomics Genomics, 783–789.10.1016/S1673-8527(07)60089-7

Gourion, B., Berrabah, F., Ratet, P., Stacey, G., 2015. Rhizobium–legume symbioses: the crucial role of plant immunity. Trends Plant Sci. 20, 186–194.10.1016/j.tplants.2014.11.008

Guerrero-Castro, J., Lozano, L., Sohlenkamp, C., 2018. Dissecting the Acid Stress Response of Rhizobium tropici CIAT 899. Front. Microbiol. 9.10.3389/fmicb.2018.00846

Guo, X., Flores, M., Mavingui, P., Fuentes, S.I., Hernández, G., Dávila, G., Palacios, R., 2003. Natural Genomic Design in Sinorhizobium meliloti : Novel Genomic Architectures. Genome Res. 13, 1810–1817.10.1101/gr.1260903

Harrison, J., Muglia, C.I., Sype, Ghislaine Van De, Aguilar, O.M., Puppo, A., Frendo, P., Jamet, A., Muglia, C.I., Van de Sype, G, Aguilar, O.M., Puppo, A., Frendo, P., 2005. Glutathione plays a fundamental role in growth and symbiotic capacity of Sinorhizobium meliloti. J Bacteriol Bacteriol, 168–174. 187/1/168[pii]10.1128/JB.187.1.168-174.2005

Herridge, D.F., Peoples, M.B., Boddey, R.M., 2008. Global inputs of biological nitrogen fixation in agricultural systems. Plant Soil Soil, 1–18.10.1007/s11104-008-9668-3

Huerta-Cepas, J., Szklarczyk, D., Heller, D., Hernández-Plaza, A., Forslund, S.K., Cook, H., Mende, D.R., Letunic, I., Rattei, T., Jensen, L.J., Von Mering, C., Bork, P., 2019. eggNOG 5.0: a hierarchical, functionally and phylogenetically annotated orthology resource based on 5090 organisms and 2502 viruses. Nucleic Acids Res. 47, D309–D314.10.1093/NAR/GKY1085

Hunter, J.D., 2007. Matplotlib: A 2D Graphics Environment. Comput. Sci. Eng. 9, 90–95.10.1109/MCSE.2007.55

Iida, T., Itakura, M., Anda, M., Sugawara, M., Isawa, T., Okubo, T., Sato, S., Chiba-Kakizaki, K., Minamisawa, K., 2015. Symbiosis Island Shuffling with Abundant Insertion Sequences in the Genomes of Extra-Slow-Growing Strains of Soybean Bradyrhizobia. Appl. Environ. Microbiol. 81, 4143–4154.10.1128/AEM.00741-15

Jenkins, C.H., Scott, A.E., O’Neill, P.A., Norville, I.H., Prior, J.L., Ireland, P.M., 2023. The Arabinose 5-Phosphate Isomerase KdsD Is Required for Virulence in Burkholderia pseudomallei. J. Bacteriol. 205.10.1128/jb.00034-23

Jones, K.M., Kobayashi, H., Davies, B.W., Taga, M.E., Walker, G.C., 2007. How rhizobial symbionts invade plants: the Sinorhizobium–Medicago model. Nat. Rev. Microbiol. 5, 619– 633.10.1038/nrmicro1705

Kalloniati, C., Tsikou, D., Lampiri, V., Fotelli, M.N., Rennenberg, H., Chatzipavlidis, I., Fasseas, C., Katinakis, P., Flemetakis, E., 2009. Characterization of a Mesorhizobium loti α-Type Carbonic Anhydrase and Its Role in Symbiotic Nitrogen Fixation. J. Bacteriol. 191, 2593–2600.10.1128/JB.01456-08

Kaneko, T., 2002. Complete Genomic Sequence of Nitrogen-fixing Symbiotic Bacterium Bradyrhizobium japonicum USDA110. DNA Res. 9, 189–197.10.1093/dnares/9.6.189

Kaneko, T., 2000. Complete Genome Structure of the Nitrogen-fixing Symbiotic Bacterium Mesorhizobium loti. DNA Res. 7, 331–338.10.1093/dnares/7.6.331

Kittell, B.L., Helinski, D.R., Ditta, G.S., 1989. Aromatic aminotransferase activity and indoleacetic acid production in Rhizobium meliloti. J. Bacteriol. 171, 5458–5466.10.1128/jb.171.10.5458-5466.1989

Kitts, P.A., Church, D.M., Thibaud-Nissen, F., Choi, J., Hem, V., Sapojnikov, V., Smith, R.G., Tatusova, T., Xiang, C., Zherikov, A., DiCuccio, M., Murphy, T.D., Pruitt, K.D., Kimchi, A., 2016. Assembly: A resource for assembled genomes at NCBI. Nucleic Acids Res. 44, D73– D80.10.1093/nar/gkv1226

Landeta, C., Davalos, A., Cevallos, M.A., Geiger, O., Brom, S., Romero, D., 2011. Plasmids with a chromosome-like role in Rhizobium. J Bacteriol. 01184-10[pii]10.1128/JB.01184-10

Liu, Y., Jiang, X., Guan, D., Zhou, W., Ma, M., Zhao, B., Cao, F., Li, L., Li, J., 2017. Transcriptional analysis of genes involved in competitive nodulation in Bradyrhizobium diazoefficiens at the presence of soybean root exudates. Sci. Rep. 7, 10946.10.1038/s41598-017-11372-0

MacLean, A.M., Finan, T.M., Sadowsky, M.J., 2007. Genomes of the symbiotic nitrogen-fixing bacteria of legumes. Plant Physiol Physiol, 615–622. 144/2/615[pii]10.1104/pp.107.101634

Mogro, E.G., Ambrosis, N.M., Lozano, M.J., 2021. Easy identification of insertion sequence mobilization events in related bacterial strains with ISCompare. G3 Genes|Genomes| Genetics 11.10.1093/G3JOURNAL/JKAB181

Nelson, M., Guhlin, J., Epstein, B., Tiffin, P., Sadowsky, M.J., 2018. The complete replicons of 16 Ensifer meliloti strains offer insights into intra-and inter-replicon gene transfer, transposon-associated loci, and repeat elements. Microb. genomics genomics, 1–11.10.1099/mgen.0.000174

Oldroyd, G.E.D., Murray, J.D., Poole, P.S., Downie, J.A., 2011. The rules of engagement in the legume-rhizobial symbiosis. Annu. Rev. Genet. 45, 119–44.10.1146/annurev-genet-110410-132549

Ormeno-Orrillo, E., Rosenblueth, M., Luyten, E., Vanderleyden, J., Martinez-Romero, E., 2008. Mutations in lipopolysaccharide biosynthetic genes impair maize rhizosphere and root colonization of Rhizobium tropici CIAT899. Env. Microbiol Microbiol, 1271–1284. EMI1541[pii]10.1111/j.1462-2920.2007.01541.

Pellock, B.J., Teplitski, M., Boinay, R.P., Bauer, W.D., Walker, G.C., 2002. A LuxR homolog controls production of symbiotically active extracellular polysaccharide II by Sinorhizobium meliloti. J. Bacteriol. 184, 5067–76.10.1128/JB.184.18.5067-5076.2002

Poole, P., Ramachandran, V., Terpolilli, J., 2018. Rhizobia: From saprophytes to endosymbionts. Nat. Rev. Microbiol. 16, 291–303.10.1038/nrmicro.2017.171

Prell, J., Bourdès, A., Karunakaran, R., Lopez-Gomez, M., Poole, P., 2009. Pathway of γ-Aminobutyrate Metabolism in Rhizobium leguminosarum 3841 and Its Role in Symbiosis. J. Bacteriol. 191, 2177–2186.10.1128/JB.01714-08

Quinlan, A.R., Hall, I.M., 2010. BEDTools: a flexible suite of utilities for comparing genomic features. Bioinformatics Bioinformatics, 841–842.10.1093/BIOINFORMATICS/BTQ033

Ruiz, B., Le Scornet, A., Sauviac, L., Rémy, A., Bruand, C., Meilhoc, E., 2019. The nitrate assimilatory pathway in Sinorhizobium meliloti: Contribution to NO production. Front. Microbiol. 10.10.3389/fmicb.2019.01526

Salas, M.E., Lozano, M.J., López, J.L., Draghi, W.O., Serrania, J., Torres Tejerizo, G.A., Albicoro, F.J., Nilsson, J.F., Pistorio, M., Del Papa, M.F., Parisi, G., Becker, A., Lagares, A., 2017. Specificity traits consistent with legume-rhizobia coevolution displayed by Ensifer meliloti rhizosphere colonization. Environ. Microbiol. 19, 3423–3438.10.1111/1462-2920.13820

Shumilina, J., Soboleva, A., Abakumov, E., Shtark, O.Y., Zhukov, V.A., Frolov, A., 2023. Signaling in Legume–Rhizobia Symbiosis. Int. J. Mol. Sci. 2023, Vol. 24, Page 17397 24, 17397.10.3390/IJMS242417397

Siguier, P., Gourbeyre, E., Chandler, M., 2014. Bacterial insertion sequences: Their genomic impact and diversity. FEMS Microbiol. Rev. 38, 865–891.10.1111/1574-6976.12067

Sobrevals, L., Müller, P., Fabra, A., Castro, S., 2006. Role of glutathione in the growth of Bradyrhizobium sp. (peanut microsymbiont) under different environmental stresses and in symbiosis with the host plant. Can. J. Microbiol. 52, 609–616.10.1139/w06-007

Stacey, G., 1995. Bradyrhizobium japonicum nodulation genetics. FEMS Microbiol. Lett. 127, 1–9.10.1111/j.1574-6968.1995.tb07441.x

Taté, R., Cermola, M., Riccio, A., Diez-Roux, G., Patriarca, E.J., 2012. Glutathione Is Required by Rhizobium etli for Glutamine Utilization and Symbiotic Effectiveness. Mol. Plant-Microbe Interact. 25, 331–340.10.1094/MPMI-06-11-0163

Tresierra-Ayala, Á., Delgado, M.J., Guzmán, R.A., Rengifo, A.L., Bedmar, E.J., 2011. MOLYBDATE TRANSPORT IN THE Bradyrhizobium japonicum - Glycine max L. SYMBIOSIS. J. soil Sci. plant Nutr. 11, 8–17.10.4067/S0718-95162011000200002

Triplett, E.W., Sadowsky, M.J., 1992. Genetics of competition for nodulation of legumes. Annu. Rev. Microbiol. 46, 399–428.10.1146/annurev.mi.46.100192.002151,

Vandecraen J., Chandler, M., Aertsen, A., Van Houdt, R., 2017. The impact of insertion sequences on bacterial genome plasticity and adaptability, Critical Reviews in Microbiology.10.1080/1040841X.2017.1303661

Virtanen, P., Gommers, R., Oliphant, T.E., Haberland, M., Reddy, T., Cournapeau, D., Burovski, E., Peterson, P., Weckesser, W., Bright, J., van der Walt, S.J., Brett, M., Wilson, J., Millman, K.J., Mayorov, N., Nelson, A.R.J., Jones, E., Kern, R., Larson, E., Carey, C.J., Polat, İ., Feng, Y., Moore, E.W., VanderPlas, J., Laxalde, D., Perktold, J., Cimrman, R., Henriksen, I., Quintero, E.A., Harris, C.R., Archibald, A.M., Ribeiro, A.H., Pedregosa, F., van Mulbregt, P., Vijaykumar, A., Bardelli, A. Pietro, Rothberg, A., Hilboll, A., Kloeckner, A., Scopatz, A., Lee, A., Rokem, A., Woods, C.N., Fulton, C., Masson, C., Häggström, C., Fitzgerald, C., Nicholson, D.A., Hagen, D.R., Pasechnik, D. V., Olivetti, E., Martin, E., Wieser, E., Silva, F., Lenders, F., Wilhelm, F., Young, G., Price, G.A., Ingold, G.L., Allen, G.E., Lee, G.R., Audren, H., Probst, I., Dietrich, J.P., Silterra, J., Webber, J.T., Slavič, J., Nothman, J., Buchner, J., Kulick, J., Schönberger, J.L., de Miranda Cardoso, J.V., Reimer, J., Harrington, J., Rodríguez, J.L.C., Nunez-Iglesias, J., Kuczynski, J., Tritz, K., Thoma, M., Newville, M., Kümmerer, M., Bolingbroke, M., Tartre, M., Pak, M., Smith, N.J., Nowaczyk, N., Shebanov, N., Pavlyk, O., Brodtkorb, P.A., Lee, P., McGibbon, R.T., Feldbauer, R., Lewis, S., Tygier, S., Sievert, S., Vigna, S., Peterson, S., More, S., Pudlik, T., Oshima, T., Pingel, T.J., Robitaille, T.P., Spura, T., Jones, T.R., Cera, T., Leslie, T., Zito, T., Krauss, T., Upadhyay, U., Halchenko, Y.O., Vázquez-Baeza, Y., 2020. SciPy 1.0: fundamental algorithms for scientific computing in Python. Nat. Methods 2020 173 17, 261–272.10.1038/s41592-019-0686-2

Walker, T.S., Bais, H.P., Grotewold, E., Vivanco, J.M., 2003. Root exudation and rhizosphere biology. Plant Physiol Physiol, 44–51.10.1104/pp.102.019661

Waskom, M.L., 2021. seaborn: statistical data visualization. J. Open Source Softw. 6, 3021.10.21105/JOSS.03021

Wheatley, R.M., Ford, B.L., Li, L., Aroney, S.T.N., Knights, H.E., Ledermann, R., East, A.K., Ramachandran, V.K., Poole, P.S., 2020. Lifestyle adaptations of Rhizobium from rhizosphere to symbiosis. Proc. Natl. Acad. Sci. U. S. A. 117, 23823–23834.10.1073/pnas.2009094117

Wheatley, R.M., Ramachandran, V.K., Geddes, B.A., Perry, B.J., Yost, C.K., Poole, P.S., 2017. Role of O 2 in the Growth of Rhizobium leguminosarum bv. viciae 3841 on Glucose and Succinate. J. Bacteriol. 199, e00572–16.10.1128/JB.00572-16

Zhang, Q., Peng, H., Gao, F., Liu, Y., Cheng, H., Thompson, J., Gao, G.F., 2009. Structural insight into the catalytic mechanism of gluconate 5-dehydrogenase from Streptococcus suis : Crystal structures of the substrate-free and quaternary complex enzymes. Protein Sci. 18, 294–303.10.1002/pro.32

